# Nicotine dosimetry and stability in Cambridge Filter PADs (CFPs) following different smoking regimen protocols and condition storage

**DOI:** 10.1101/2020.09.09.289371

**Authors:** Pietro Zuccarello, Sonja Rust, Massimo Caruso, Rosalia Emma, Roberta Pulvirenti, Claudia Favara, Riccardo Polosa, Giovanni Li Volti, Margherita Ferrante

**Affiliations:** Department of Medical, Surgical Sciences and Advanced Technologies “G.F. Ingrassia”, Via S. Sofia, 87, 95123 Catania (Italy); Center of Excellence for the Acceleration of Harm Reduction (CoEAHR), University of Catania, Via S. Sofia, 97, 95123, Catania (Italy); Department of Biomedical and Biotechnological Sciences, Via S. Sofia, 97, 95123 Catania (Italy); Department of Clinical and Experimental Medicine, University of Catania, Via S. Sofia, 97, 95123, Catania (Italy)

**Keywords:** nicotine, electronic cigarette, stability, Cambridge Filter Pad, smoking regimen, storage

## Abstract

Despite the growing numbers of studies with cigarettes and other electronic nicotine delivery products (ENDs), there is no standard covering nicotine dosimetry and its stability in various matrix. The aim of the present study was to provide a protocol to normalize nicotine concentration adsorbed in Cambridge Filter PADs (CFPs) and their storage method. Smoke/vapor generated by a reference tobacco cigarette (1R6F) and ENDs with different exposure regimes (ISO, HCI and CRM81) was collected in CFPs. For each exposure, some CFPs were analyzed at time zero, whereas the others were stored under different conditions for nicotine assessment after 30 days. Principal Component Analysis (PCA) was also performed to establish the best parameter for nicotine normalization. PCA showed the best correlation between nicotine in CFPs and TPM. Our results showed differences between products and puffing regimes, but storage of CFPs at −80°C was always effective in maintaining the nicotine content. In conclusion, this study highlights that different exposure regimens and products can affect the preservation of nicotine titer in CFPs and samples storage at −80°C may prevent the loss of nicotine. These conditions are recommended and should be adopted for Inter-laboratory comparison studies on ENDs to ensure harmonization between participating laboratories.

## INTRODUCTION

Nicotine is an alkaloid extracted from tobacco leaves. It is a dibasic compound with pyridine and pyrrolidine rings and a pKa of 8.5. Nicotine is water soluble and separates preferentially by organic solvents depending on the solution pH. Its degradation mechanisms include photolysis, thermolysis, oxidation, and hydrolysis (1, 2). In fact, it is a colorless photosensitive substance being easily oxidized when exposed to light or air, turning to a brown color (3). Therefore, pharmaceutical formulations containing nicotine must be stored in the dark and at temperatures not exceeding 25 °C (4). Moreover, some environmental bacteria and fungi may be also responsible for nicotine degradation (5).

Nicotine exhibits a wide spectrum of toxicological profile also related to its thermal degradation metabolites. Indeed, following heating, nicotine is degraded producing nitrogen oxides, carbon monoxide and other highly toxic fumes. Therefore, over the last decade consumer products able to deliver nicotine by a combustion-free process has significantly increased. These products, known as electronic cigarettes (e-cigs) and tobacco heating products (THPs) seem to be a less harmful alternative to smoking because provide ‘smoking experience without smoking’ (6–8).

The potential benefits and risks of using combustion-free nicotine delivery technologies, (e-cig and THP) also known as Reduced Risk Products (RRPs) have been the subject of intense scientific debate (9). *In vitro* studies allow to carry out a quick and easy evaluation of the potential human health impact of these products. A number of toxicological tests is required to establish the reduced harm potential compared to combustible cigarettes and to ensure protection of individual and public health from the adverse effects of potentially harmful exposures (9, 10).

In order to clarify the toxicological effects of RRPs compared to combustible cigarette, confirmatory and, possibly, multicenter studies are warranted. Therefore, standardized protocols allowing to reproduce *in vitro assays* are warranted in order to obtain consistent results in terms of biological effect.

Various smoking regimens and approaches have been developed and implemented to measure smoke constituents of tobacco products including the International Organization of Standardization (ISO)(11) and Health Canada Intensive (HCI) smoking regimens (12). Each of these smoking regimens has been developed for different reasons. ISO regimen is considered a non-intense smoking regimen otherwise HCI is considered an intense smoking regimen. The different intensities of the smoking regimens are used to understand and quantify the various levels of harmful constituents to which consumers may be exposed (13). Since 2000 the Health Canada Intense (HCI) regimen with hole vents blocked was used to collect cigarette smoke components and, recently, for the collection of aerosol components from the Tobacco Heating Products (THPs). However, for these latter products, the hole vents are not-blocked in order to avoid the overheating of electronic devices. During 2014 the CORESTA E-cigarette Task Force drafted a Technical Report outlining the necessary requirements for the generation and collection of e-cigarette aerosol for analytical testing purposes and defined the exposure regimen for e-cigarettes, the "CORESTA Reference Method n.81" (CRM81) (14). Moreover, although tobacco cigarette and RRPs are often considered as similar products, they exhibit several differences regarding released substances during their operation. Tobacco products and RRPs electronic devices share similar release of nicotine in smoke/aerosols. Thus, the comparison between these two types of products is often made on the basis of the nicotine released during their use. Therefore, it is relevant to establish a standard for nicotine dosimetry and to determine its stability in order to schedule *in vitro* experiments accordingly. However, there is no data available on the stability of nicotine to other recommended stressors or smoking regimen.

The aims of the present study were at first to evaluate the best way to normalize the nicotine concentration adsorbed in Cambridge Filter PADs (CFPs) during exposure, then to conduct comparisons between the different storing conditions for the nicotine in CFPs.

## METHODS

### Cambridge Filter Pads (CFP) exposure

Three products were used to expose CFPs: 1R6F reference cigarettes (University of Kentucky), Vype ePen3 and Vype eStick Maxx electronic cigarettes (Nicoventures Trading Ltd). Borgwaldt LM1 smoking machine and LM4E vaping machines (Borgwaldt KC, Hamburg – Germany) were used to collect total particulate matter (TPM) from smoke and vapor respectively (Figure 1).

**Figure 1.**
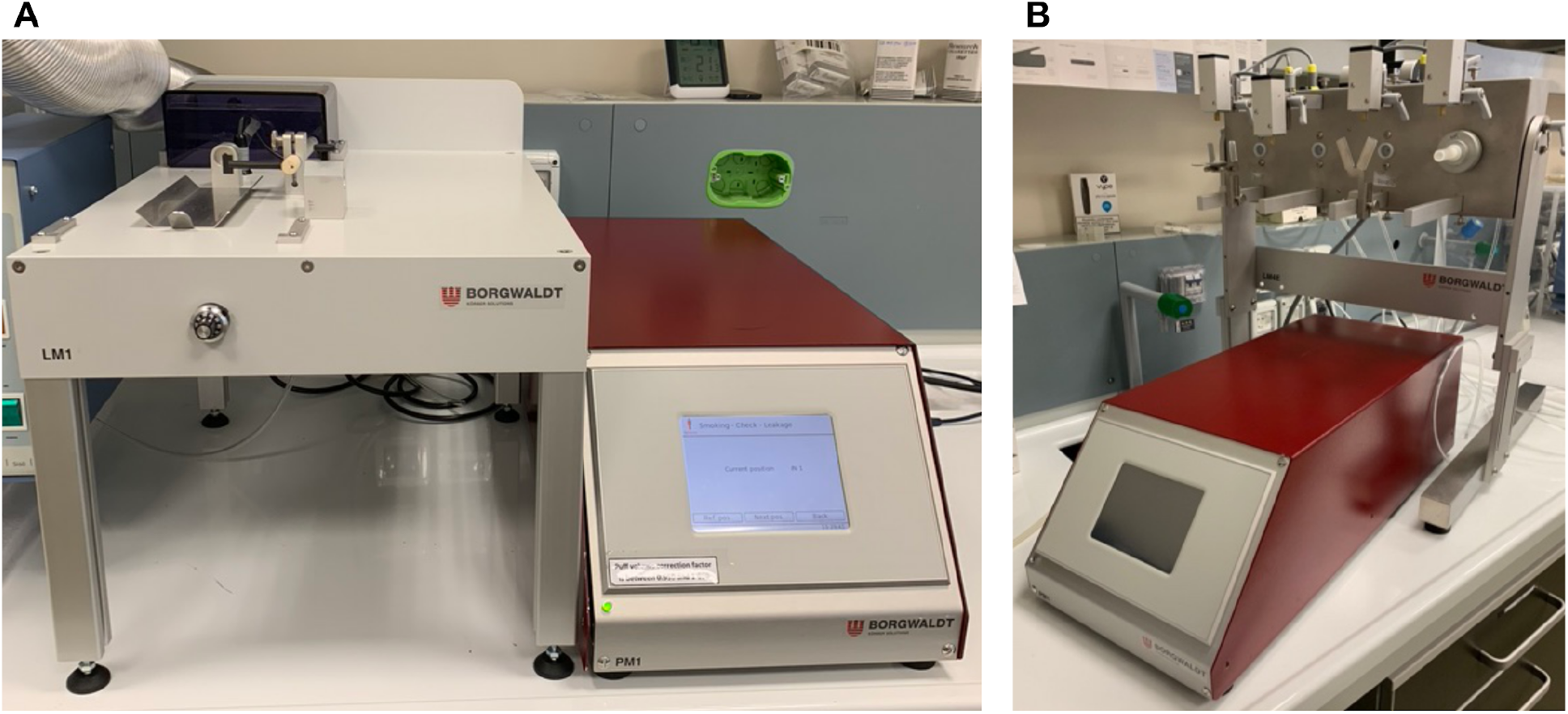
The Borgwaldt LM1 (A) smoking machine and LM4E button automated (B) vaping machine. Undiluted aerosol is generated from a single syringe (within the red box) and delivered from devices into the corresponding in vitro exposure system.

The 1R6F reference cigarettes are unflavored blended cigarettes (83 mm length) characterized by 0.721 mg/cigarette of nicotine following International Organization for Standardization (ISO) smoking regime, and 1,896 mg/cigarette of nicotine following Health Canada Intense (HCI) smoking regimen, as reported by Kentucky University (15). Vype ePen3 is a button-activated “closed-modular” e-cigarette, while Vype eStick Maxx is a puff-activated cigarette-like product. Both devices consist of two modules, a rechargeable battery section and a replaceable liquid (“e-liquid”) containing cartridge (“cartomizer”). “Master Blend” flavored variant containing 18 mg/mL nicotine was used for Vype ePen3, and “Toasted Tobacco” flavored variant containing 18 mg/mL nicotine was used for Vype eStick Maxx.

1R6F cigarettes were conditioned for at least 48h (60 ± 3% relative humidity, 22 ± 1 °C) before smoke generation, and smoked in a test atmosphere of 60 ± 5% relative humidity, 22 ± 2 °C according to ISO 3402:1999. Nineteen reference cigarettes 1R6F were smoked to the length of the filter +8 mm at ISO regimen (35 mL puff volume, drawn over 2 s, once every minute with ventilation holes unblocked) according to reference method described in ISO 4387:2000, and nineteen reference cigarette 1R6F were smoked following the HCI regimen (55 mL puff volume, drawn over 2 s, once every 30 s with ventilation holes blocked) to the length of the filter +8mm. The smoke generated by each cigarette was captured in line on a 44 mm diameter CFP for each one. Vype ePen3 was vaped following a modified HCI regimen (55 mL puff volume, drawn over 2 s, once every 30 s with square shape profile) plus 1 s of pre-activation, for 15 puffs/CFP. Vype eStick Maxx was vaped following CRM81 regimen (55 mL puff volume, drawn over 3 s, once every 30 s with square shape profile) for 15 puffs/CFP.

### Sample preparation

Sample preparation and analysis was performed according with the international standard (16). After collection of TPM (Total Particulate Matter), each Cambridge Filter Pad (CFP) was chopped into small pieces and transferred into a 15 mL plastic tube containing 10 mL of extraction solvent consisting of isopropanol (LC/MS grade, Carlo Erba) with N-decane (purity 99%, Sigma-Aldrich) (50 μg/mL) as internal standard. Tubes were shaked for 30 minutes by vortex at 200 rpm. The samples were then sonicated for 5 minutes in an ultrasonication bath. Subsequently, 1 ml of each sample were filtered with cellulose acetate filters (mm 25; um 0.45) and 100 μl of each extract were transferred in a vial with a conical insert for auto-sampler.

### GC-FID analysis

Analysis were performed by a gas chromatography Shimadzu (model GC 2010 AF) coupled with Flame Ionization Detector. An Agilent J&W DB-HeavyWAX Intuvo GC column (30 m × 0.25 mm, 0.25 μm) was used. The GC-FID operating condition and the column oven temperature program are reported in table 1 and table 2, respectively.

**Table 1.** GC-FID operating condition.

|  |  |  |
| --- | --- | --- |
| Injection | Temperature | 250°C |
|  | Mode | Split |
|  | Volume | 1 µl |
| Carrier Gas: H2 | Flow control mode | Pressure |
|  | Pressure | 68 kPa |
|  | Flow | 40 ml/min |
| Make Up Gas: N2 | Flow | 30 ml/min |
| Air | Flow | 400 ml/min |
| Detector | Temperature | 250 °C |
|  | Sampling rate | 40msec |

**Table 2.** Column oven temperature program.

|  | Rate °C/min | Temperature °C | Hold time min |
| --- | --- | --- | --- |
| 0 | - | 50 | 0 |
| 1 | 60 | 170 | 5 |
| 2 | 80 | 250 | 1 |

### Calibration curve

Nicotine stock solution at concentration of 100 μg/μL was prepared weighing 1 g of nicotine at purity of 99% (Sigma Aldrich) into a 10 mL volumetric flask and dilute to volume with acetone. The solution was stored between 0 °C and 4 °C in dark. Nicotine calibrating standard solutions were prepared at concentration levels 0, 100, 200, 500 and 1000 μg/mL in 1 milliliter of extraction solution consisting to propan-2-ol with heptadecane at purity of 99% (Sigma Aldrich, cod. 128503-100G) at concentration of 50 μg/L.

### Dosimetry performance assessment

For linearity assessment, calibration curve was performed, and linear correlation coefficient was assessed (r^2^): the acceptability criterion was r^2^> 0.98. For the precision and accuracy assessment, eight filters CFP were spiked with nicotine at concentration of 100 μg for CFP. Precision was assessed on the basis of the Relative Standard Deviation (RSD%) as the percental ratio between Standard Deviation and the mean value: the acceptability criterion was RSD%<10%. The accuracy was estimated on the base of the Recovery (R%) as the percent ratio between the main value and the real dose (100 μg): the acceptability criterion was R%>80%.

### Nicotine normalization assessment

The Principal Component Analysis (PCA) was used to verify the correlation of variables (concentration of nicotine on filters, TPM and cartridge weight gap or number of puff) and determine if the normalization of the nicotine concentration on filters for TPM was relevant, according to CORESTA indication. We performed one PCA for electronic devices (Vype ePen and eStick) and one for the two regimens of exposure (ISO and HCI) for reference cigarettes 1R6F.

The Principal Components Analysis (PCA) was performed using RStudio software Version 1.2.5033. The regularized discriminant analysis (RDA) function was used for PCA. Data were standardized before analysis and the results were displayed in a biplot of correlation. The filter samples were unscaled and the weighted dispersion was equal on all dimensions. The variables were scaled proportionally to eigenvalues. Although the small number of variables, the sample size was considered appropriate for this purpose (17).

### Comparison between the different storing conditions

RStudio Software was also used to perform the statistical analysis aimed to assess stability of nicotine in different storing conditions. For both each product and exposure regime data were collected in five groups based on the storage conditions (Tables 3, 4, 5 and 6). Three CFPs, analyzed at zero time without any conditioning, for each product/exposure regime, were included into the control groups (group 0). For each product/exposure regime, 4 filters were stored for 30 days after exposure, under different conditions: (i) at room temperature (25 °C) in extraction solution (group 1), (ii) at room temperature (25 °C) dry (group 2), (iii) at the temperature of −20 °C dry (group 3) and, finally, (iv) at the temperature of −80 °C dry (group 4). Therefore, the CFPs from all four groups (1, 2, 3 and 4) for each storage condition were analyzed for nicotine content. For each CFP, the concentration of nicotine was normalized for the weight of total particulate matter (TPM) both for reference cigarettes and for electronical devices.

**Table 3.** Data collected for each CFP exposed to vapor from Vype ePen and for each storing condition about nicotine concentration, Total Particulate Matter (TPM) and Cartridge Weight (CW) CFPs.

| **CFP – Storing condition groups** | **Nicotine**<br>**(µg/CFP)** | **TPM (mg)** | **CW (mg)** |
| --- | --- | --- | --- |
| Control<br>(0) | 898 | 78.1 | 93.5 |
|  | 944 | 79.1 | 95.2 |
|  | 1102 | 92.3 | 117.6 |
| Extraction solution<br>(1) | 1013 | 85.1 | 100.5 |
|  | 1000 | 82.6 | 100.3 |
|  | 1008 | 81.0 | 96.5 |
|  | 970 | 80.3 | 95.3 |
| Room Temperature (25 °C)<br>(2) | 862 | 83.6 | 101.9 |
|  | 832 | 75.7 | 90.1 |
|  | 917 | 79.5 | 98.0 |
|  | 983 | 84.4 | 99.4 |
| Temperature = -20 °C<br>(3) | 1029 | 89.2 | 107.3 |
|  | 887 | 77.3 | 94.5 |
|  | 1006 | 80.9 | 99.5 |
|  | 957 | 79.6 | 94.4 |
| Temperature = -80 °C<br>(4) | 1128 | 95.6 | 121.2 |
|  | 931 | 85.6 | 97.3 |
|  | 993 | 79.9 | 95.4 |
|  | 1032 | 82.4 | 99.2 |
| **Mean Value** | **973.2** | **82.7** | **99.8** |
| **SD** | **75.9** | **5.1** | **7.8** |
| **RSD%** | **7.8** | **6.2** | **7.9** |

**Table 4.** Data collected for each CFP exposed to vapor from Vype eStick and for each storing condition about nicotine concentration, Total Particulate Matter (TPM) and Cartridge Weight (CW).

| **CFP – Storing condition groups** | **Nicotine**<br>**(µg/CFP)** | **TPM (mg)** | **CW (mg)** |
| --- | --- | --- | --- |
| Control<br>(0) | 567 | 62.5 | 68.0 |
|  | 557 | 62.1 | 67.8 |
|  | 499 | 54.8 | 58.3 |
| Extraction solution<br>(1) | 565 | 56.6 | 107.3* |
|  | 432 | 50.8 | 50.5 |
|  | 321 | 36.7 | 41.0 |
|  | 423 | 46.4 | 52.4 |
| Room Temperature (25 °C)<br>(2) | 382 | 41.6 | 47.6 |
|  | 347 | 42.9 | 44.6 |
|  | 387 | 44.4 | 47.6 |
|  | 374 | 45.1 | 47.8 |
| Temperature = -20 °C<br>(3) | 525 | 55.0 | 60.7 |
|  | 469 | 50.6 | 54.8 |
|  | 387 | 42.3 | 47.0 |
|  | 372 | 40.8 | 46.7 |
| Temperature = -80 °C<br>(4) | 429 | 49.6 | 53.2 |
|  | 417 | 46.1 | 48.1 |
|  | 426 | 43.7 | 50.0 |
|  | 373 | 40.0 | 45.6 |
| **Mean Value** | **434.3** | **48.0** | **51.8** |
| **SD** | **75.6** | **7.4** | **7.6** |
| **RSD%** | **17.4** | **15.4** | **14.6** |
| * anomalous value excluded for determination of mean value, SD and RSD% |  |  |  |

**Table 5.** Data collected for each CFP exposed to smoke from 1R6F reference cigarette by ISO regimen and for each storing condition about nicotine concentration, number of puffs and Total Particulate Matter (TPM).

| **CFP – Storing condition groups** | **Nicotine**<br>**($\mu\text{g}/\text{CFP}$)** | **Number of puffs** | **TPM mg** |
| --- | --- | --- | --- |
| Control<br>(0) | 441 | 6.7 | 7.6 |
|  | 492 | 6.0 | 7.7 |
|  | 465 | 7.0 | 7.7 |
| Extraction solution<br>(1) | 458 | 6.0 | 7.7 |
|  | 533 | 6.0 | 8.5 |
|  | 467 | 7.0 | 7.7 |
|  | 455 | 7.2 | 7.3 |
| Room Temperature (25 °C)<br>(2) | 350 | 6.7 | 8.0 |
|  | 402 | 6.8 | 8.5 |
|  | 262 | 6.6 | 7.7 |
|  | 400 | 7.0 | 8.3 |
| Temperature = -20 °C<br>(3) | 366 | 6.2 | 7.6 |
|  | 340 | 6.4 | 7.0 |
|  | 393 | 7.3 | 8.0 |
|  | 472 | 6.3 | 8.1 |
| Temperature = -80 °C<br>(4) | 399 | 6.0 | 8.0 |
|  | 559 | 6.3 | 7.7 |
|  | 388 | 7.5 | 7.9 |
|  | 572 | 6.6 | 8.6 |
| **Mean Value** | **432.4** | **6.6** | **7.9** |
| **SD** | **78.0** | **0.5** | **0.4** |
| **RSD%** | **18.0** | **7.1** | **5.2** |

**Table 6.** Data collected for each CFP exposed to smoke from 1R6F reference cigarette by HCI regimen and for each storing condition about nicotine concentration, number of puffs and Total Particulate Matter (TPM).

| **CFP – Storing condition groups** | **Nicotine**<br>**($\mu\text{g}/\text{CFP}$)** | **Number of puffs** | **TPM mg** |
| --- | --- | --- | --- |
| Control<br>(0) | 1366 | 9.2 | 29.3 |
|  | 1287 | 7.7 | 30.1 |
|  | 1245 | 8.2 | 28.6 |
| Extraction solution<br>(1) | 1368 | 8.0 | 35.0 |
|  | 1294 | 8.0 | 32.2 |
|  | 1441 | 7.3 | 29.6 |
|  | 1421 | 8.3 | 31.2 |
| Room Temperature<br>(2) | 1179 | 8.0 | 26.6 |
|  | 1234 | 7.4 | 27.4 |
|  | 1178 | 8.7 | 29.9 |
|  | 1289 | 7.6 | 32.2 |
| Temperature = -20 °C<br>(3) | 1205 | 8.2 | 29.4 |
|  | 1410 | 7.9 | 31.4 |
|  | 1493 | 8.0 | 32.7 |
|  | 1379 | 8.4 | 30.2 |
| Temperature = -80 °C<br>(4) | 1473 | 9.0 | 34.5 |
|  | 1507 | 8.0 | 33.2 |
|  | 1295 | 8.4 | 29.7 |
|  | 1415 | 8.4 | 31.1 |
| **Mean Value** | **1341.0** | **8.1** | **30.8** |
| **SD** | **105.3** | **0.5** | **2.2** |
| **RSD%** | **7.8** | **6.0** | **7.2** |

For each group, normality was assessed by Shapiro-Wilk test. A p-value >0.05 was considered for a normal distribution. Moreover, for each group, a one-sample t-test was performed to assess the homogeneity of the average value of the group with the average value of the control group. A p-value >0.05 was considered for the homogeneity of the average values.

Finally, the non-parametric test of Kruskal-Wallis was performed to investigate the differences of the storing conditions compared to the control group. Dunn’s-test for multiple comparisons of independent samples was used as post-hoc test. A p-value <0.05 was for a statistically significative difference.

## RESULTS

### Dosimetry performance assessment

The linear correlation coefficient r^2^ of calibration curve was 0.9999. Method showed at concentration level of 100 μg/filter an RSD% and a recovery equal to 6.0% and 94.6%, respectively.

### Vape/Smoke exposure in CFP

For each exposure run with vapor from electronic cigarettes (ePen and eStick), data on weight of both cartridge and CFPs, before and after the puff run, were collected. For each exposure run with smoke from 1R6F reference cigarettes, both with ISO 3308:2000 and HCI regimens, data on weight of CFPs before and after the puff run were collected. The difference between the weight of the CFP after and before the exposure to smoke/vapor is considered as the total particulate matter (TPM). Data collected from CFPs exposed to e-cigs vapor are shown in tables 3 and 4. Data on CFPs exposed to smoke from 1R6F reference cigarette, both with ISO 3308:2000 and HCI regimens, are shown respectively in Tables 5 and 6. The means ± SD weights of TPM collected from ePen and eStick are 87.7 ± 5.1 mg and 48.0 ± 7.8, respectively. Whereas, the means ± SD weights of TPM collected from 1R6F following ISO and HCI regimes are 6.6 ± 0.5 mg and 8.1 ± 0.5 mg.

### Nicotine normalization assessment

Correlation biplot for ePen (PC1 96% and PC2 2%) showed a high correlation between nicotine concentration on filters and TPM (Figure 2). PCA highlight a minor correlation with the cartridge weight gap. Samples between 1 and 19 are relative to ePen exposure while samples between 20 and 38 are relative to eStick exposure. The two group of exposure are located in two different area.

**Figure 2.**
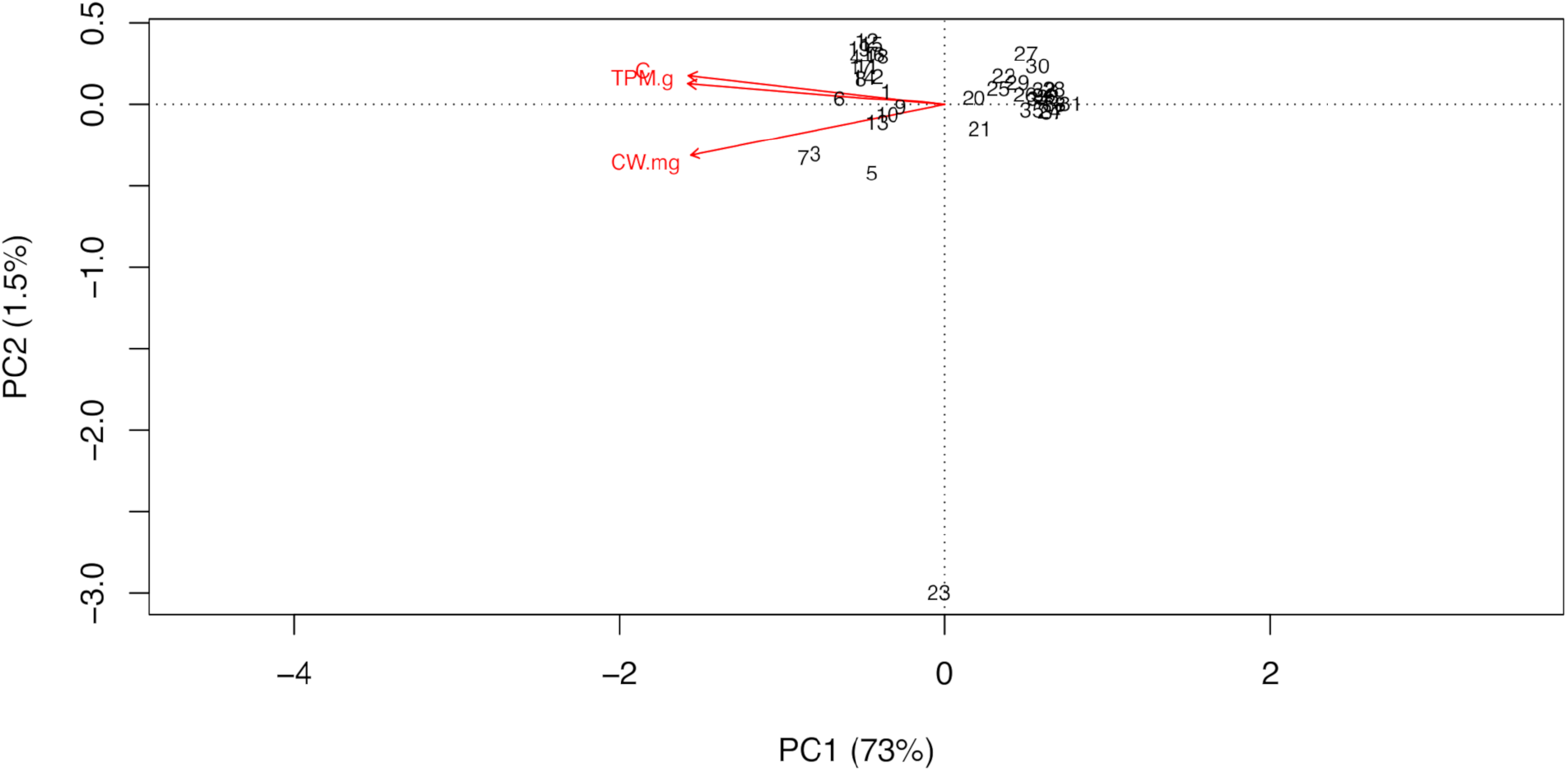
Correlation biplot for e-cigarettes: samples between 1 and 19 are relative to ePen exposure, while samples between 20 and 38 are relative to eStick exposure.

Correlation biplot for 1R6F (PC1 96% and PC2 2%) shows also a high correlation between nicotine concentration on filters and TPM. PCA highlight a minor correlation with the number of puffs. Samples between 1 and 19 are relative to ISO-regime exposure while samples between 20 and 38 are relative to HCI-regime exposure. The two group of exposure regime are located in two different longitudinal bands (Figure 3).

**Figure 3.**
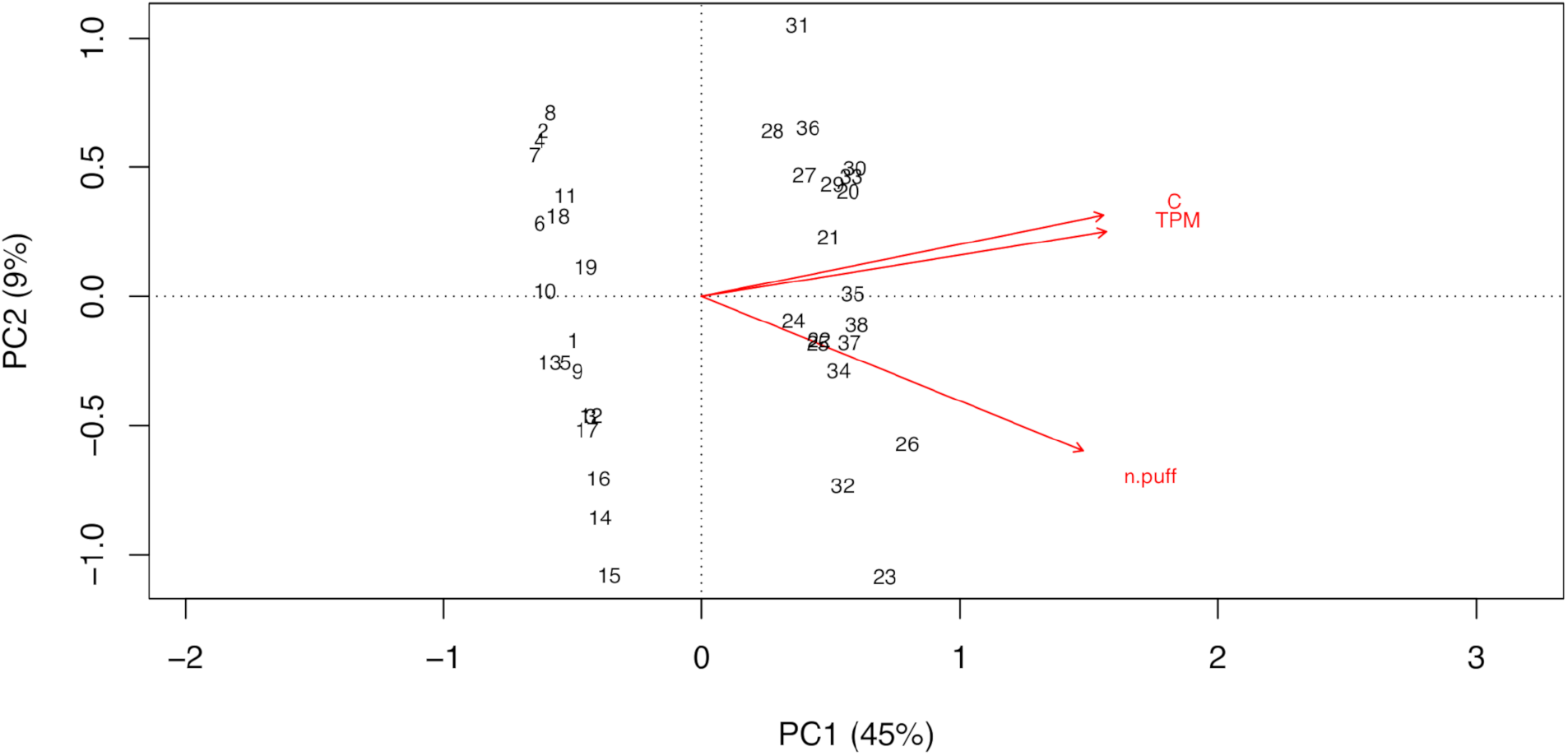
Correlation biplot for 1R6F: samples between 1 and 19 are relative to ISO-regime exposure while samples between 20 and 38 are relative to HCI-regime exposure.

### Comparison between the different storing conditions

Data of nicotine concentrations normalized for TPM are reported in table 7. For Vype ePen, distributions of nicotine normalized for TPM of the groups 1, 2, 3 and 4 were normal (p-values were 0.5719 for the group 1, 0.6298 for the group 2, 0.2742 for the group 3 and 0.3741 for the group 4). Indeed, the normality for the control group was not verified (p-value was 2.2E-16).

**Table 7.** Nicotine content normalized to TPM for each CFP exposed to vapor from electronic cigarettes (Vype ePen and eStick) and smoke from 1R6F reference cigarette by HCI and ISO regimens and for each storing condition.

| **TPM-normalized Nicotine** | **ePen** | **eStick** | **ISO regimen** | **HCI regimen** |
| --- | --- | --- | --- | --- |
| **Control**<br>(0) | 11.5 | 9.1 | 58.0 | 46.6 |
|  | 11.9 | 9.0 | 63.9 | 42.8 |
|  | 11.9 | 9.1 | 60.4 | 43.5 |
| **Extraction solution**<br>(1) | 11.9 | 10.0 | 59.5 | 39.1 |
|  | 12.1 | 8.5 | 62.7 | 40.2 |
|  | 12.4 | 8.7 | 60.6 | 48.7 |
|  | 12.1 | 9.1 | 62.3 | 45.6 |
| **Room Temperature**<br>(2) | 10.3 | 9.2 | 43.8 | 44.3 |
|  | 11.0 | 8.1 | 47.3 | 45.0 |
|  | 11.5 | 8.7 | 34.0 | 39.4 |
|  | 11.7 | 8.3 | 48.2 | 40.0 |
| **Temperature = -20 °C**<br>(3) | 11.5 | 9.5 | 48.2 | 41.0 |
|  | 11.5 | 9.3 | 48.6 | 44.9 |
|  | 12.4 | 9.2 | 49.1 | 45.7 |
|  | 12.0 | 9.1 | 58.3 | 45.7 |
| **Temperature = -80 °C**<br>(4) | 11.8 | 8.6 | 49.9 | 42.7 |
|  | 10.9 | 9.0 | 72.6 | 45.4 |
|  | 12.4 | 9.8 | 49.2 | 43.6 |
|  | 12.5 | 9.3 | 66.5 | 45.5 |
| **Mean Value** | **11.8** | **9.0** | **54.9** | **43.7** |
| **SD** | **0.6** | **0.5** | **9.4** | **2.7** |
| **RSD%** | **4.9** | **5.2** | **17.1** | **6.1** |
| * anomalous value excluded from statistical analysis |  |  |  |  |

Homogeneity of the mean value of the control group and the mean value of groups 2, 3 and 4 was verified (p-values were 0.1319 for the group 2, 0.7277 for the group 3 and 0.7408 for the group 4). Indeed, for 1^st^ group homogeneity was not verified (p-values were 0.04016). Kruskal-Wallis test did not show a statistically significative difference between the storing conditions and the control group (p-value was 0.1439) (Figure 4).

**Figure 4.**
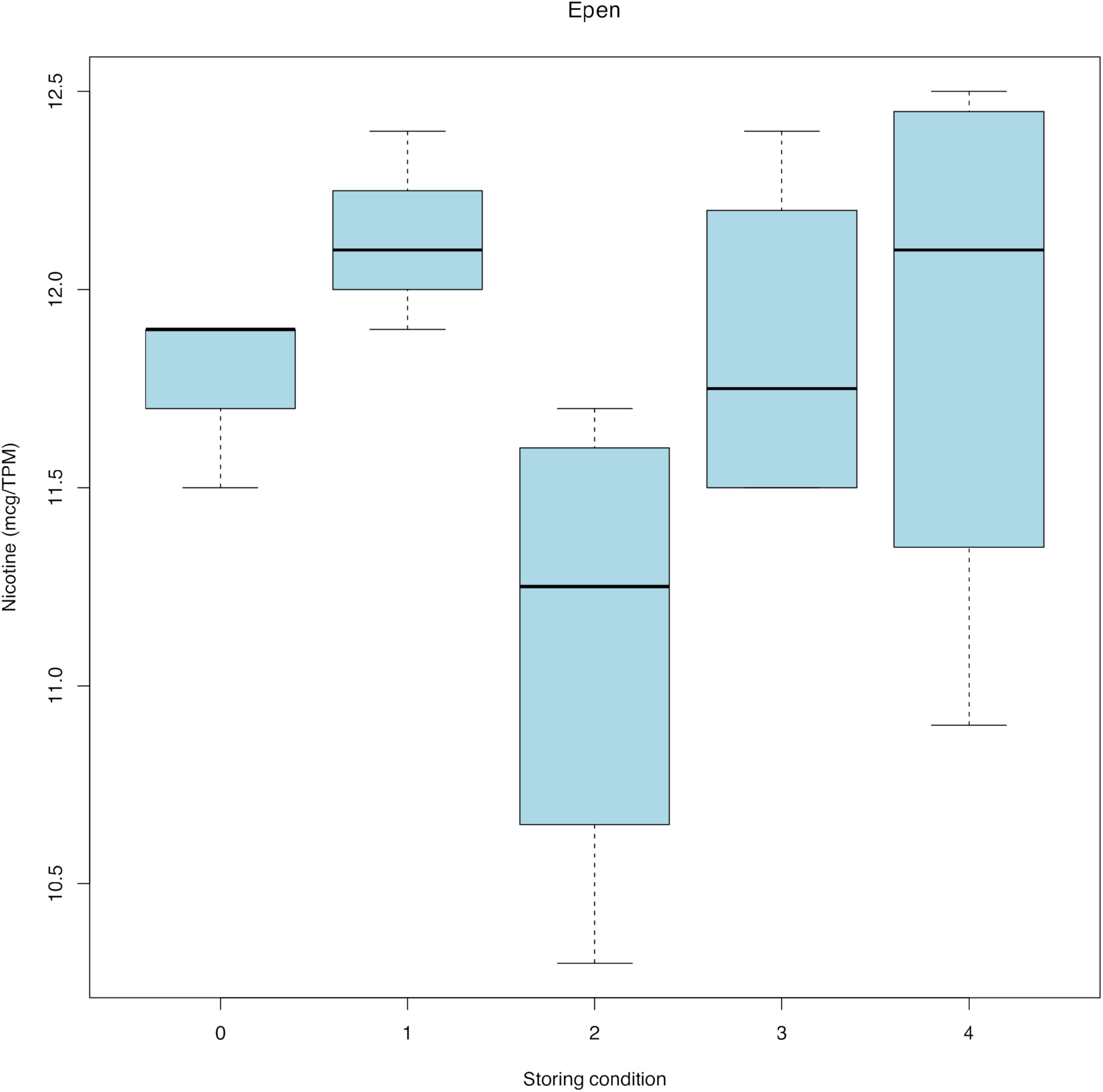
Box plot for exposure of Vype e-Pen between the TPM-normalized nicotine and the storing conditions. The mean value and the standard deviation were respectively 11.8 and 0.23 mcg/TPM for control group (group 0), 12.1 and 0.21 mcg/TPM for group 1, 11.1 and 0.63 mcg/TPM for the group 2, 11.9 and 0.44 mcg/TPM for the group 3 and, finally, 11.9 and 0.73 mcg/TPM for the group 4.

For eStick, distributions of nicotine normalized for TPM of the groups 1, 2, 3 and 4 were normal (p-values were 0.4679 for the group 1, 0.7553 for the group 2, 0.85 for the group 3 and 0.9886 for the group 4). The normality for the control group was not verified (p-value was 2.2E-16). Homogeneity of the mean value of each group and the mean value of the control group was verified (p-values were 0.9816 for the group 1, 0.136 for the group 2, 0.09252 for the group 3 and, finally, 0.6973 for the group 4). Kruskal-Wallis test did not show a statistically significative difference between the storing conditions and the control group (p-value was 0.2851) (Figure 5).

**Figure 5.**
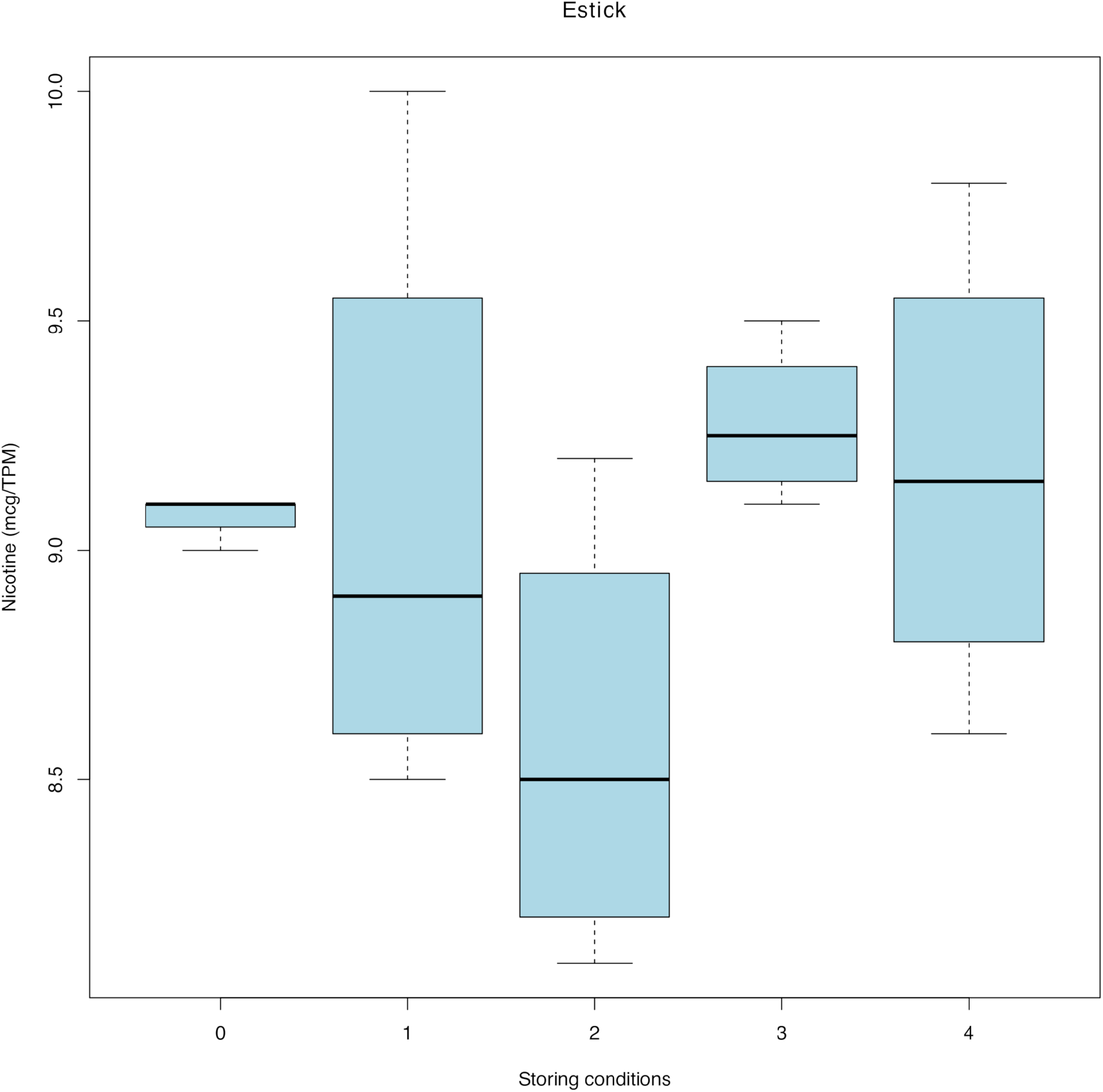
Box plot for exposure of Vype eStick between the TPM-normalized nicotine and the storing conditions. The mean value and the standard deviation were respectively 9.1 and 0.06 mcg/TPM for control group (group 0), 9.1 and 0.67 mcg/TPM for group 1, 8.6 and 0.49 mcg/TPM for the group 2, 9.3 and 0.17 mcg/TPM for the group 3 and, finally, 9.2 and 0.51 mcg/TPM for the group 4.

For reference cigarettes 1R6F exposed in ISO regimen, distributions of nicotine normalized for TPM of control group and groups 1, 2 and 4 were normal (the p values were 0.7952 for control group, 0.5436 for the group 1, 0.2066 for the group 2 and, finally, 0.2188 for the group 4). The normality for the group 3 was not verified (p=0.01162).

Homogeneity of the mean value of the groups 1 and 4 and the mean value of the control group was verified (p-values were 0.5448 for the group 1 and 0.85 for the group 4). It was not verified of for groups 2 and 3 (p-values were 0.01266 for the group 2, 0.02784 for the group 3).

Kruskal-Wallis showed a statistically significative difference between the storing conditions and the control group (p-value was 0.01226) (Figure 6). Post-hoc Dunn’s test show only a statistically significative difference between the group 2 and the control group (p=0.0127).

**Figure 6.**
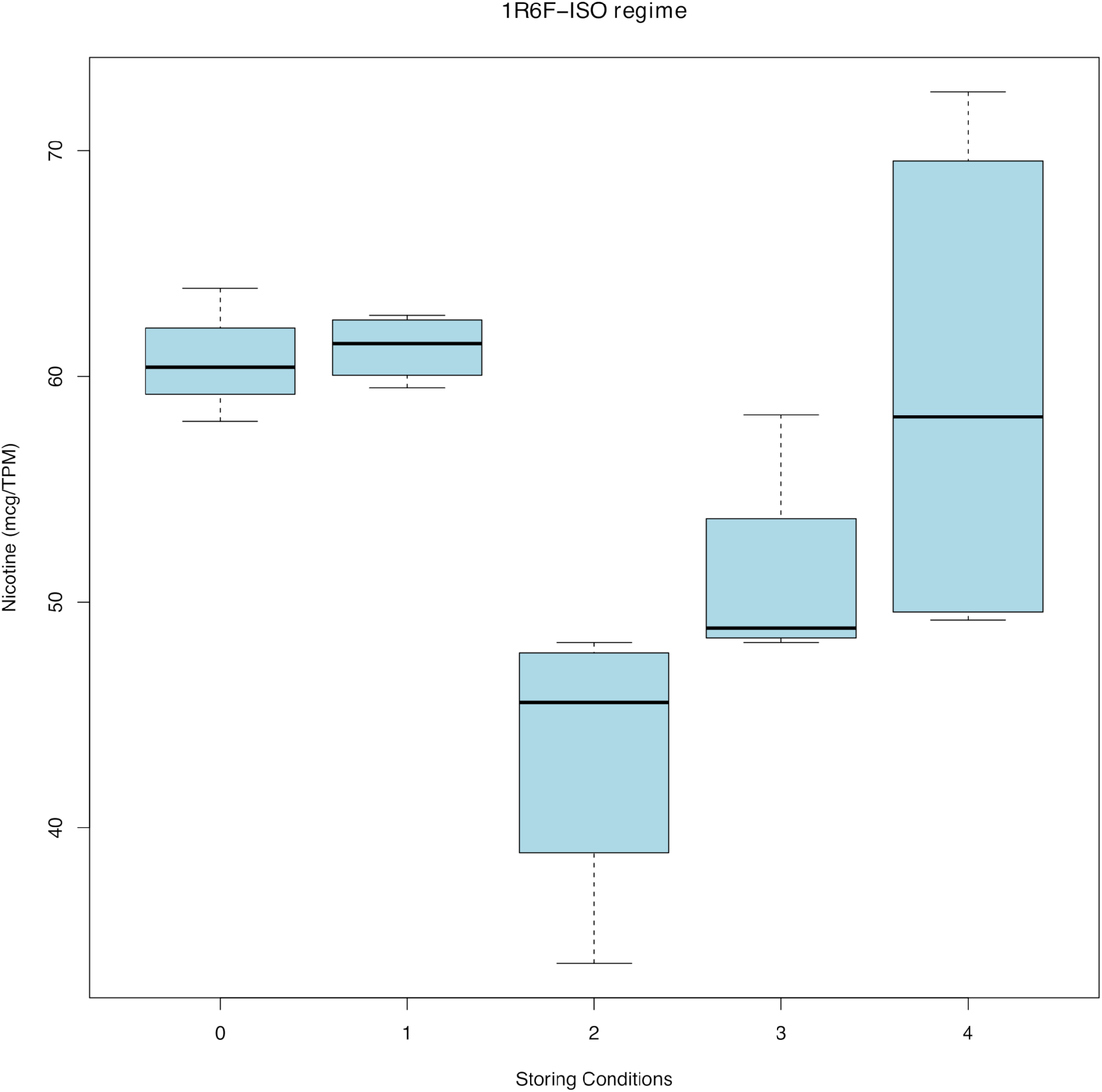
Box plot for exposure of 1R6F in ISO regimen between the TPM-normalized nicotine and the storing conditions. The average value and the standard deviation were respectively 60.8 and 2.96 mcg/TPM for control group (group 0), 61.3 and 1.49 mcg/TPM for group 1, 43.3 and 6.50 mcg/TPM for the group 2, 51.1 and 4.85 mcg/TPM for the group 3 and, finally, 59.6 and 11.8 mcg/TPM for the group 4.

For reference cigarettes 1R6F exposed in HCI regimen, distributions of nicotine normalized for TPM of control group and the group 1, 2 and 3 were normal (p-values were 0.3322 for control group, 0.4892 for the group 1, 0.1829 for the group 2 and 0.2882 for the group 4). Indeed, the normality for the group 3 was not verified (p=0.03522).

Homogeneity of the mean value of each group and the mean value of the control group was verified (p-values were 0.7179 for the group 1, 0.2368 for the group 2, 0.9837 for the group 3 and, finally, 1.0000 for the group 4). Kruskal-Wallis test did not show a statistically significative difference between the storing conditions and the control group (p-value was 0.7156) (Figure 7).

**Figure 7.**
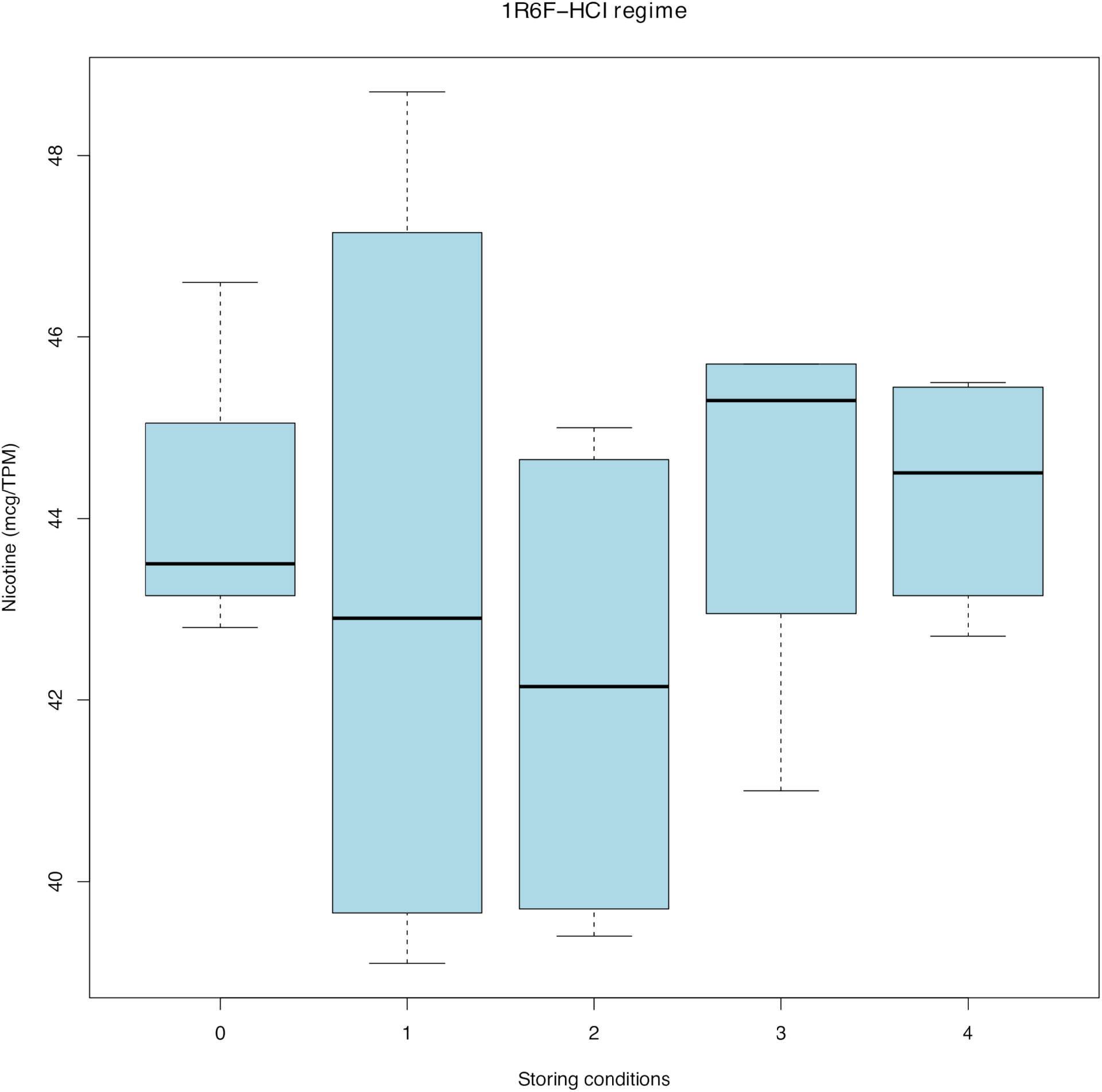
Box plot for exposure of 1R6F in HCI regime between the TPM-normalized nicotine and the storing conditions. The mean value and the standard deviation were respectively 44.3 and 2.02 mcg/TPM for control group (group 0), 43.4 and 4.53 mcg/TPM for group 1, 42.2 and 2.88 mcg/TPM for the group 2, 44.3 and 2.25 mcg/TPM for the group 3 and, finally, 44.3 and 1.38 mcg/TPM for the group 4.

## DISCUSSION

Nicotine dosimetry, for the purpose of comparing exposure from different products (cigarettes, e-cig, THP), is a common and often used practice in studies that aim to evaluate the different impact of cigarette smoke and vapors produced by alternative products on different cell systems (18–22). The nicotine retention and its stability in the CFPs over time is a fundamental validation step for smoke particulate phase science. The use of CFP to collect and analyze TPM constituents was first referenced approximately 50 years ago (Wartman et al., 1959). Previously, the stability of total particulate matter retained on CFPs was evaluated by assessing its ability to induce genotoxicity and cytotoxicity *in vitro*, but none of its components, including nicotine, has been determined (23, 24). A standardized and reliable nicotine dosimetry test applied to various electronic devices and combustible cigarettes is therefore of great importance when evaluating the accuracy of the exposures carried out in *in vitro* assays and its impact on cellular response when using the same exposure conditions (21).

Furthermore, since the smoker who is switching to electronic devices gets satisfaction only upon reaching the same blood concentration of nicotine acquired with combustion cigarette (25), it is necessary to compare between the number of puffs of tobacco products of combustible cigarette and electronic nicotine devices required to achieve the same dose of nicotine.

Such practice is also of particular importance when performing Ring Studies with several participants where it is necessary to perform nicotine dosimetry before *in vitro* treatments in order to obtain reliable and comparable results among all partners. Indeed, there is no certainty that the exposure performed in one partner’s laboratory will be comparable to the ones performed in other satellite laboratories if standards are not in place. For this reason, it also needs to be taken into account that nicotine stability during shipment are an intrinsic part of Nicotine dosimetry analysis. Time, temperature, humidity, light and air exposure are factors to keep in account during storage and transit. In a Ring study these tests are usually conducted by one leading lab who receives the samples by its international partners (26).

To this regard, no protocols on nicotine samples storage (i.e. filter PADs), shipping temperature or extraction solvent are so far available. To assess these conditions, we used nicotine as a reference since it is a major substance released from both smoke and vapor from all products. Then we measured it in the CFPs to evaluate whether the same exposure and the same number of puffs, at different regimen produces the same quantity and quality of smoke/vapor. This is an important step to harmonize protocols prior to conducting any in vitro assays. Many studies did not address issues relating to storage and stability of the CFPs. Since Watson and colleagues noted the semi volatility and degradability of nicotine (27), data on both the loss rate for nicotine and the stability under different storage conditions, such as room temperature, refrigerated (4° C), frozen (−20° C), and ultralow freezing (−70° C) would be desirable.

Furthermore, our results showed no significant differences of nicotine stability when using Vype eStick or reference cigarette in HCI regimen (p>0.05). Interestingly, when using HCI regimen, we observed no significant difference in nicotine concentration in all the experimental groups when compared to control, whereas when using ISO regimen, we observed a significant difference of mean value in CFP group stored at room temperature and at temperature −20°C (p<0.05). Moreover, a significant difference of variance in CFP group stored at room temperature was also shown.

Such difference in nicotine stability between the two smoking regimens may be dependent on different oxygenation obtained with the two protocols due to the entry of air from the cigarette filter holes in the ISO regimen (free filter holes) compared to the HCI regimen (blocked filter holes) leading to increased nicotine oxidation with the first regimen.

Regarding the exposure by ePen, a significant difference (p<0.05) was shown for the mean value of the CFPs group stored in solvent solution, higher than the mean value of CFPs of the control group. The long contact time of the CFPs with the extraction solvent could have significantly improved the extraction efficiency and increased the nicotine concentrations compared to the extemporaneous extraction of the samples.

However, although it was not always statistically significant, the nicotine concentrations found in CFPs stored at room temperature was always lower than that of the control group and the other groups. This denotes a sensitive degradation of nicotine even at temperatures below 30 °C.

On the other hand, the preservation of the samples at −80 °C has always proved effective in maintaining the nicotine content in CFPs.

The Principal Component Analysis showed a high correlation between nicotine and TPM values. This shows that the delivery of nicotine is directly proportional to that of the particulate matter. In terms of RSD%, the exposure with ISO regime showed a high inaccuracy in the release of TPM (15.4%) and the number of puffs (14.6%), as well as of the nicotine concentrations found in CFPs (17.4%), although conserved under different conditions. The normalization of the nicotine concentrations in the CFPs for the amount of TPM released during the exposure allowed to reduce the variability of the study due to the precision and accuracy of the exposure process by the smoking and the vaping machine. By this way it was possible to believe that the nicotine values in the CFPs were homogeneous at the time of exposure and that the change was due to the method and the time of storage of the samples.

## CONCLUSION

In conclusion, this study highlights that different exposure regimens and different products can affect the preservation of nicotine titer in CFPs. However, refrigerating the samples at minus 80 °C up to 30 days before analysis prevents the oxidative and thermal degradation effects on the substance. This storing procedure of CFPs storing is strongly recommended for the standardization of protocol and it should be required for Ring studies on tobacco and nicotine containing products to be used in exposure systems.

## REFERENCES

1. Benowitz, N.L., Hukkanen, J. and Jacob, P. (2009) Nicotine chemistry, metabolism, kinetics and biomarkers. Handbook of experimental pharmacology, 2009: 10.1007/978-3-540-69248-5_2.

2. George Ngwa (2010) Forced Degradation as an Integral Part of HPLC Stability-Indicating Method Development. Drug Deliv. Technol, 10, 56–59.

3. Mishra, A., Chaturvedi, P., Datta, S., Sinukumar, S., Joshi, P. and Garg, A. (2015) Harmful effects of nicotine. Indian journal of medical and paediatric oncology : official journal of Indian Society of Medical & Paediatric Oncology, 36, 24–31.

4. American Society of Health System Pharmacists (2009) AHFS Drug Information 2009. Bethesda, MD, 2009.

5. Brandsch, R. (2006) Microbiology and biochemistry of nicotine degradation. Applied microbiology and biotechnology, 69, 493–8.

6. Caponnetto, P., Russo, C., Bruno, C.M., Alamo, A., Amaradio, M.D. and Polosa, R. (2013) Electronic cigarette: a possible substitute for cigarette dependence. Monaldi archives for chest disease = Archivio Monaldi per le malattie del torace, 79, 12–9.

7. Farsalinos, K.E., Romagna, G., Tsiapras, D., Kyrzopoulos, S. and Voudris, V. (2014) Characteristics, perceived side effects and benefits of electronic cigarette use: a worldwide survey of more than 19,000 consumers. International journal of environmental research and public health, 11, 4356–73.

8. Biener, L. and Hargraves, J.L. (2015) A longitudinal study of electronic cigarette use among a population-based sample of adult smokers: association with smoking cessation and motivation to quit. Nicotine & tobacco research : official journal of the Society for Research on Nicotine and Tobacco, 17, 127–33.

9. Polosa, R., O’Leary, R., Tashkin, D., Emma, R. and Caruso, M. (2019) The effect of e-cigarette aerosol emissions on respiratory health: a narrative review. Expert review of respiratory medicine, 13, 899–915.

10. Bals, R., Boyd, J., Esposito, S., Foronjy, R., Hiemstra, P.S., Jiménez-Ruiz, C.A., et al. (2019) Electronic cigarettes: A task force report from the European Respiratory Society. European Respiratory Journal, 53.

11. ISO - ISO 3308:2012 - Routine analytical cigarette-smoking machine — Definitions and standard conditions. https://www.iso.org/standard/60404.html (30 July 2020).

12. ISO/TR 19478-2:2015(en), ISO and Health Canada intense smoking parameters — Part 2: Examination of factors contributing to variability in the routine measurement of TPM, water and NFDPM smoke yields of cigarettes. https://www.iso.org/obp/ui/#iso:std:iso:tr:19478:-2:ed-1:v1:en (30 July 2020).

13. Marian, C., O’Connor, R.J., Djordjevic, M. V., Rees, V.W., Hatsukami, D.K. and Shields, P.G. (2009) Reconciling human smoking behavior and machine smoking patterns: Implications for understanding smoking behavior and the impact on laboratory studies. Cancer Epidemiology Biomarkers and Prevention, 18, 3305–3320.

14. CORESTA RECOMMENDED METHOD N^o^ 81 - ROUTINE ANALYTICAL MACHINE FOR E-CIGARETTE AEROSOL GENERATION AND COLLECTION – DEFINITIONS AND STANDARD CONDITIONS (2015) 2015. https://www.coresta.org/sites/default/files/technical_documents/main/CRM_81.pdf (27 July 2020).

15. Center for Tobacco Reference Products - University of Kentucky (2018) Certificate of Analysis-1R6F Certified Reference Cigarette. 2018. https://ctrp.uky.edu/assets/pdf/webdocs/CoA18_1R6F.pdf (27 July 2020).

16. No. 7 - Determination of Nicotine in the Mainstream Smoke of Cigarettes by Gas Chromatographic Analysis | CORESTA. https://www.coresta.org/determination-nicotine-mainstream-smoke-cigarettes-gas-chromatographic-analysis-29141.html (30 July 2020).

17. Zuccarello, P., Ferrante, M., Cristaldi, A., Copat, C., Grasso, A., Sangregorio, D., et al. (2019) Reply for comment on &Exposure to microplastics (. Water research, 166, 115077.

18. Bode, L.M., Roewer, K., Otte, S., Wieczorek, R. and Simms, L. (2017) Dosimetry: The effects of cigarette smoke dilution on nicotine delivery in a 24 and 96 well format using Smoke Aerosol Exposure In Vitro System (SAEIVS). CORESTA Meeting, Smoke Science/Product Technology, 2017, Kitzbühel, STPOST 22, 2017. www.imperialbrandsscience.com (30 July 2020).

19. Wieczorek, R.L., Trelles Sticken, E., Bode, L.M. and Simms, L. (2017) Dosimetry: TPM and nicotine delivery of combusted tobacco products to the 24 and 96 Multi Well Plate and inserts on Smoke Aerosol Exposure In vitro System (SAEIVS). CORESTA Meeting, Smoke Science/Product Technology, 2017, Kitzbühel, STPOST 23, 2017. www.imperialbrandsscience.com (30 July 2020).

20. Adamson, J., Thorne, D., Zainuddin, B., Baxter, A., McAughey, J. and Gaça, M. (2016) Application of dosimetry tools for the assessment of e-cigarette aerosol and cigarette smoke generated on two different in vitro exposure systems. Chemistry Central Journal, 10, 74.

21. Adamson, J., Li, X., Cui, H., Thorne, D., Xie, F. and Gaca, M.D. (2017) Nicotine Quantification In Vitro : A Consistent Dosimetry Marker for e-Cigarette Aerosol and Cigarette Smoke Generation. Applied In Vitro Toxicology, 3, 14–27.

22. Behrsing, H., Hill, E., Raabe, H., Tice, R., Fitzpatrick, S., Devlin, R., et al. (2017) In Vitro Exposure Systems and Dosimetry Assessment Tools for Inhaled Tobacco Products: Workshop Proceedings, Conclusions and Paths Forward for *In Vitro* Model Use. Alternatives to Laboratory Animals, 45, 117–158.

23. Wartman, W.B., Cogbill, E.C. and Harlow, E.S. (1959) Determination of Particulate Matter in Concentrated Aerosols: Application to Analysis of Cigarette Smoke. Analytical Chemistry, 31, 1705–1709.

24. Crooks, I., Dillon, D.M., Scott, J.K., Ballantyne, M. and Meredith, C. (2013) The effect of long term storage on tobacco smoke particulate matter in in vitro genotoxicity and cytotoxicity assays. Regulatory Toxicology and Pharmacology, 65, 196–200.

25. Marsot, A. and Simon, N. (2016) Nicotine and Cotinine Levels with Electronic Cigarette. International Journal of Toxicology, 35, 179–185.

26. Garner, C.R.J., Reynolds, U.S.A., Stevens, R.D. and Tayyarah, R. (2015) E-Cigarette Task Force Technical Report 2014 Electronic Cigarette Aerosol Parameters Study.

27. Watson, C., McCraw, J., Polzin, G., Ashley, D. and Barr, D. (2004) Development of a method to assess cigarette smoke intake. Environmental science & technology, 38, 248–53.

